# Non-invasive Differentiation of M1 and M2 Activation in Macrophages using Hyperpolarized ^13^C MRS of Pyruvate and DHA at 1.47 Tesla

**DOI:** 10.1101/2020.12.02.405845

**Authors:** Kai Qiao, Lydia M. Le Page, Myriam M. Chaumeil

**Author notes:** **Corresponding author:** Myriam M. Chaumeil, Ph.D., 1700 4th Street, BH 204, Mission Bay Campus Box 2530, San Francisco, CA 94143.

## Abstract

Macrophage activation, first generalized to the M1/M2 dichotomy, is a complex and central process of the innate immune response. Simply, M1 describes the classical pro-inflammatory activation, leading to tissue damage, and M2 the alternative activation promoting tissue repair. Given the central role of macrophages in multiple diseases, the ability to non-invasively differentiate between M1 and M2 activation states would be highly valuable for monitoring disease progression and therapeutic responses. Since M1/M2 activation patterns are associated with differential metabolic reprogramming, we hypothesized that hyperpolarized ^13^C magnetic resonance spectroscopy (HP ^13^C MRS), an innovative metabolic imaging approach, could distinguish between macrophage activation states noninvasively. The metabolic conversions of HP [1-^13^C]pyruvate to HP [1-^13^C]lactate and HP [1-^13^C]dehydroascorbic acid to HP [1-^13^C]ascorbic acid were monitored in live M1 and M2 activated J774a.1 macrophages non-invasively by HP ^13^C MRS on a 1.47 Tesla NMR system. Our results show that both metabolic conversions were significantly increased in M1 macrophages compared to M2 and non-activated cells. Biochemical assays and high resolution ^1^H MRS were also performed to investigate the underlying changes in enzymatic activities and metabolite levels linked to M1/M2 activation. Altogether, our results demonstrate the potential of HP ^13^C MRS for monitoring macrophage activation states non-invasively.

## Introduction

Nearly ubiquitous throughout the body, macrophages are a critical component in maintaining our health and wellbeing, playing a central role in the innate immune response, tissue homeostasis, and facilitation of crosstalk with neighboring cell types^1-3^. Macrophages have typically been described as having two activation states, which have each been observed to play significant roles in various pathologies, such as autoimmune disorders, obesity, and cancer malignancy^4-6^. Although there is still much to be understood, these activation states were first generalized to the M1 and M2 dichotomy^7^, and are associated with cell-wide changes, including modulations of signaling pathways and reprogramming of cellular metabolism^8^. Briefly, M1 describes the classical pro-inflammatory activation response, leading to subsequent tissue damage, whereas M2 activation is associated with upregulation of anti-inflammatory pathways promoting tissue repair. From an energetic metabolism perspective, M1 macrophages have been shown to increase anaerobic glycolysis, while M2 use primarily aerobic oxidative phosphorylation for energy generation^9^. Reactive oxygen species (ROS) have also been reported to increase significantly in M1 activated macrophages and are often used as marker of this activation state^10^. The activity of the arginase-1 enzyme isoform meanwhile has been shown to significantly increase specifically in M2 activated macrophages, serving as a robust marker of M2 activation^11^. Thus far, M1 and M2 differentiation has been largely determined using such markers by histological, biochemical, and genotypic studies, conducted on samples collected invasively from the bloodstream or diseased tissues^12^. To date no *in vivo* imaging method enables differentiation between M1 and M2 activation states. Given their broad and critical roles, such an approach could be very useful in improving our understanding of the role of macrophages in underlying diseases, and would also allow *in vivo* monitoring of response to immunomodulatory treatments targeting macrophages activation *in situ*.

An investigative tool for assessment of *in vivo* metabolism exists, in the form of ^13^C magnetic resonance spectroscopy (MRS) combined with dissolution dynamic nuclear polarization (dDNP) of ^13^C-labeled probes^13^, so called hyperpolarized (HP) ^13^C MRS. This technique has been used to noninvasively monitor metabolic impairment in multiple preclinical models, including cancer, neuroinflammation, and cardiomyopathy^14-16^. Importantly, the use of this technology is now expanding into clinical settings, with ongoing clinical trials on brain tumor patients, traumatic brain injury patients, and prostate cancer patients, amongst others^17-19^. Depending on the choice of ^13^C labeled substrate, different metabolic pathways can be targeted and imaged non-invasively^20,21^. To date, two studies have used HP ^13^C MRS to look at activation states of macrophages. One study described the used of HP [6-^13^C]arginine to detect arginase activity in M2-like primary mouse myeloid-derived suppressor cells (MDSCs)^22^. The authors showed that metabolism of HP [6-^13^C]arginine to HP ^13^C urea by arginase was significantly increased in MDSCs compared to control bone marrow cells, in line with increased arginase activity linked to M2 activation. Another study on J774a.1 macrophage cells using HP [1-^13^C]pyruvate showed that, after M1 activation using the toxin lipopolysaccharides (LPS), the conversion of HP [1-^13^C]pyruvate into [1-^13^C]lactate catalyzed by the lactate dehydrogenase (LDH) enzyme was significantly increased compared to non-treated macrophages. This result was linked to, amongst other cellular events, increased LDH activity and gene transcription^23^. However, to date, no study has directly compared M1 vs. M2 activation states in macrophages using HP ^13^C MRS.

Here, we described a comprehensive study in which we applied HP ^13^C MRS to paired M1 vs. control and paired M2 vs. control macrophages. Two metabolic conversions were evaluated: 1) the conversion of HP [1-^13^C]pyruvate into HP [1-^13^C]lactate, and 2) the conversion of HP [1-^13^C]dehydroascorbic acid (DHA) into HP [1-^13^C]ascorbic acid (AA) (**Figure 1A**). The first reaction is the final step of glycolysis, while the second reaction is modulated by the levels of ROS. Both reactions have previously been successfully imaged *in vivo*^24,25^. In our study, we show that for both HP probes, HP product-to-substrate ratios were observed to be significantly increased in M1 macrophages compared to M2, while control and M2 macrophages presented similar metabolic ratios. Underlying mechanisms were investigated, and spectrophotometric assays showed that changes in HP lactate-to-pyruvate ratios were associated with changes in LDH and pyruvate dehydrogenase (PDH) enzymatic activities, while changes in HP AA-to-DHA ratio paralleled changes in ROS levels.

**Figure 1.**
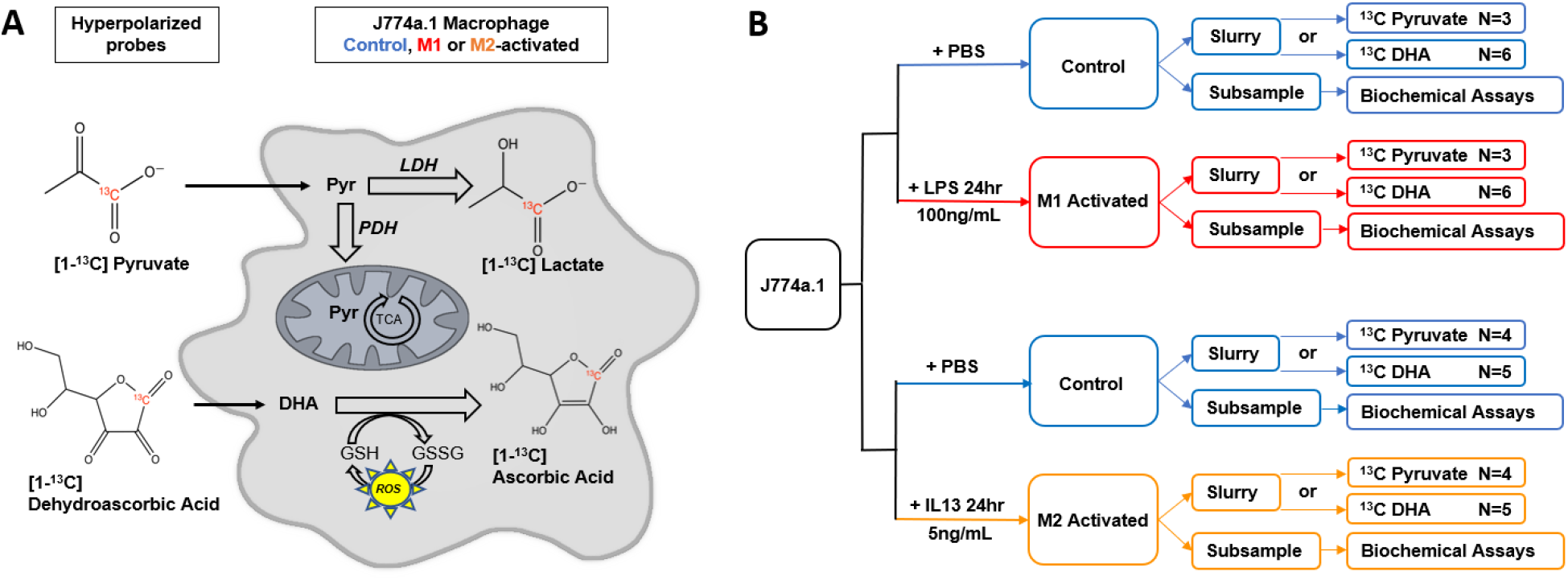
Overall design. (**A**) Schematic of hyperpolarized ^13^C probes and their metabolic fates in J774a.1 macrophages. LDH=lactate dehydrogenase; PDH=pyruvate dehydrogenase; TCA=tricarboxylic acid cycle; GSH=glutathione; GSSG=glutathione disulfide; ROS=reactive oxygen species. (**B**) Overall experimental design of the study. Activation was achieved with either lipopolysaccharide (LPS, M1 activation) or interleukin-13 (IL-13, M2 activation). Control and M1/M2 activated samples were paired for HP studies and for spectrophotometric assays.

Our study therefore shows that noninvasive differentiation of M1 vs M2 activation states in macrophages can be achieved using ^13^C MRS of HP [1-^13^C]pyruvate and HP [1-^13^C]DHA at the clinically relevant field strength of 1.47 Tesla, demonstrating the potential of this method to evaluate immune response and to monitor effect of immunomodulatory treatments.

## Material and Methods

### Cell Culture

J774a.1 mouse macrophages (ATCC) were grown in Dulbecco’s Modified Eagle’s Media (DMEM) containing 10% fetal bovine serum and 5% Penicillin/Streptomycin (UCSF). M1 activation was achieved with 100ng/mL lipopolysaccharide (LPS) treatment for 24h (*E. coli*; Sigma Aldrich), and M2 activation with 5ng/mL of murine interleukin-13 (IL-13) for 24h (Peprotech), as previously described^23,26^. A control group was established with a vehicle treatment of sterile PBS. All passages numbers used were ~4 – 14 to reduce the risk of genetic drift, and a mycoplasma testing kit (ATCC) confirmed the culture was contamination-free. **Figure 1B** presents a schematic of the full experimental design.

For HP ^13^C experiments, cells were studied as paired samples of either M1 or M2 activation and paired controls. The definition of paired samples follows: Three T225 flasks were split into ten T225 flasks, five of which were activated towards either M1 or M2, and the other five with vehicle as control. The adherent macrophages were incubated with 0.04% EDTA in calcium and magnesium-free PBS solution (Cell Culture Facility, UCSF) for approximately 10 minutes, collected and then centrifuged at 125xG for 5 minutes. The pellet was resuspended in 200μL of fresh DMEM (no additives), and a 20μL sample was taken and washed with PBS to be saved at −80°C for paired spectrophotometric assays (for all assays except ROS). Another 20μL sample was taken for cell counting (referred to as subsample in the rest of this manuscript), and the remaining slurry suspension (~20 million cells) was transferred to a 5mm NMR tube. All injections of HP probes were done within 5 minutes of cell resuspension and transfer to the NMR tube.

### MR Acquisitions

24μL of [1-^13^C]pyruvate (15M pyruvic acid (Sigma), 15mM Trityl radical (GE), 500mM Gd-DOTA (Guerbet)) or 25μL of [1-^13^C]DHA (2.2M dehydroascorbic acid (Sigma) prepared as previously described^25^) was polarized for 1h on a Hypersense dDNP polarizer (Oxford Instruments), then dissolved in 4.5mL or 3.5mL buffer (pyruvate buffer: 80mM NaOH, 40mM Tris HCl, 3mM EDTA in ddH_2_O; DHA buffer: 3mM EDTA in ddH_2_O), to yield a final solution of 80mM or 15.72mM, respectively. Within 20s of dissolution, approximately 400μL of HP [1-^13^C]pyruvate (n = 3 Control vs. M1-activated pairs, n = 4 Control vs. M2-activated pairs) or [1-^13^C]DHA (DHA: n = 6 Control vs. M1 activated pairs, n = 5 Control vs. M2 activated pairs) was injected into a 5mm NMR tube containing a ~20 million cells slurry in 200uL DMEM. Hyperpolarized spectra were then acquired on a 1.47T Oxford Pulsar NMR system using the following parameters: Flip angle = 20°, repetition time (TR) = 3s, and number of scans (NS) = 100 for a total acquisition of 5 minutes. Analysis was performed with Mestrenova (Mestrelab) software. Signal to noise ratio (SNR) for the injected substrates and detected products were calculated from the total summed spectra as area under curve divided by standard deviation of the noise. All HP data are represented as mean ± standard error of the mean (SEM) and are normalized to cell number and volume of injection.

### Spectrophotometric Assays

ROS levels, arginase activity, LDH activity, PDH activity, and glutathione (GSH) levels were measured with spectrophotometry on paired samples collected from the same flasks as the ones used for HP experiments. ROS data of control (n = 6), M1 activated (n = 6), and M2 activated (n = 3) was reported as fold-change from control using a commercial intracellular ROS fluorescence assay kit (Abcam, used according to manufacturer’s instructions). Arginase assay (Abcam) activity of control (n = 6), M1 activated (n = 3), and M2-activated (n = 3) was reported in units/Liter (U/L) and normalized to 10^5^ cells per well. An in-house LDH assay measuring the rate of NADH (Sigma) depletion for control (n = 6) and activated groups (n = 3 for both M1 and M2) was performed, and reported as μM NADH/minute normalized to protein concentration quantified by Bradford assay (Thermofisher). A PDH assay kit (Abcam) was used to measure PDH enzyme activity between groups (n = 3 for each control, M1, and M2), and reported as optical density (milliOD)/minute with normalization to protein concentration. GSH and its oxidized form glutathione disulfide (GSSG) were measured with a commercial kit (Biovision). The total glutathione levels were first measured, then subtracted from measured levels of GSH to obtain GSSG values, with the subsequent GSH/GSSG ratio reported with normalization to 10^5^ cells per well. All spectrophotometric data are reported as mean ± standard deviation.

### High resolution ^1^H NMR of extracted cell metabolites

Metabolites from M1 activated (n = 5), M2 activated (n = 5), and Control (n = 4) J774a.1 cells were extracted using equal parts methanol-water-chloroform, as previously described^27^. Cold, 4°C saline (5mL) was added directly to T75 flasks 2-3 times and removed to rinse, and −20°C methanol (3mL) was subsequently added. The adherent macrophages were scraped off into the methanol – this mixture was transferred to a clean tube. Equal parts −20°C chloroform and 4°C H_2_O were homogenously mixed, and the fractions separated by centrifuging at 125xG at 4°C. The methanol fraction was collected, 11.7mM Trimethylsilylpropanoic acid (TSP) (Acros Organics) added, and the mixture lyophilized. The resultant extracts were reconstituted in 420uL D_2_O, and the samples were scanned on an 800Mhz NMR system (Bruker) with a 1D NOESY sequence and NS = 384. Spectral processing was performed with Mestrenova (Mestrelab), and a select group of metabolites of interest previously reported in macrophage studies^23^ were fitted and quantified using Chenomx NMR Suite (Chenomx Inc) with reference to the Human Metabolomics Database^28^. Concentrations of quantified metabolites were normalized to cell number and TSP reference, and reported as mean ± standard deviation.

### Statistical analyses

The sample sizes of HP experiments were determined using an 80% power calculation of preliminary paired data between activated and non-activated groups. All data were tested with two-way ANOVA between activated and non-activated, with Sidak’s multiple comparisons test. All tests were performed with Prism 8 (GraphPad) software.

## Results

### M1 and M2 activation of J774a.1 macrophages

First, ROS and Arginase assays were performed to confirm differential M1 and M2 activation of macrophages. Arginase enzyme activity was unchanged with M1 activation (p = 0.849), while a significant 57-fold increase in Arginase activity was seen in M2 activated macrophages (p < 0.0001, n = 3, 38.255 U/L/10^5^ cells in M2 activated *vs*. 0.669 U/L/10^5^ cells for Control), as previously reported^26^ (**Figure 2A**). On the other hand, M1 activated macrophages exhibited a 2.3-fold increase in ROS compared to Control (p = 0.0009, n = 6), whereas no significant increase in ROS was detected in M2 activated macrophages (p = 0.7380, n = 3) (**Figure 2B**). The ROS increase in M1 activated macrophages is in agreement with previous reports^29^. The Bradford assay showed no differences in total protein concentration per cell between groups (**Figure 2C**).

**Figure 2.**
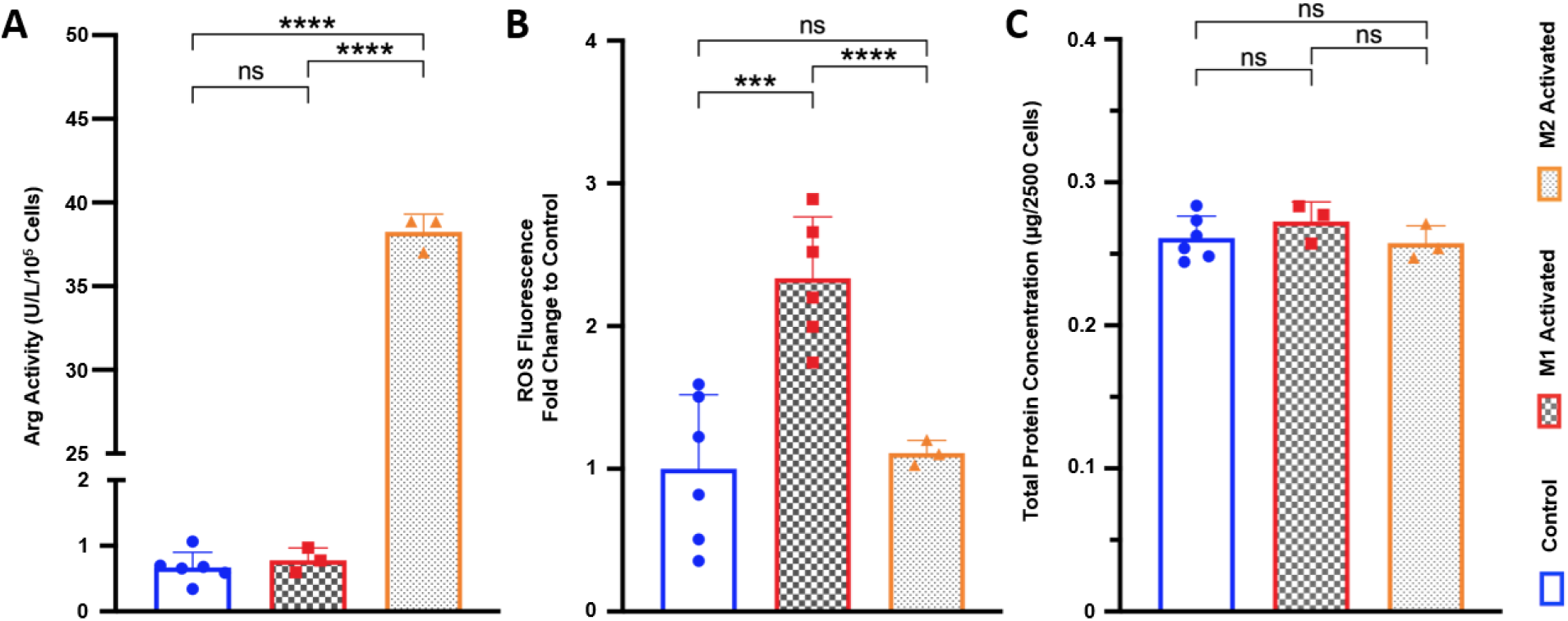
M1 and M2 activation of J774a.1 macrophages. (**A**) Arginase (Arg) activity in Units (where 1 Unit = amount of arginase that will generate 1.0 nmol of H_2_O_2_/min at 37°C) for control (blue), M1 activated (red) and M2 activated (orange) macrophages (ns: p = 0.849, ****p<0.0001). (**B**) Reactive oxygen species (ROS) fluorescence levels between control (blue), M1 activated, (red) and M2 activated (orange) J774a.1 macrophages, expressed in fold change compared to control (ns: p = 0.738, ***p<0.001, ****p<0.0001). (**C**) Total protein concentrations normalized to cell numbers for control (blue), M1 activated, (red) and M2 activated (orange) J774a.1 macrophages (ns control vs M1: p = 0.309; ns control vs M2: p = 0.721; ns M1 vs M2: p = 0.229).

### HP [1-^13^C]lactate production is differentially increased by M1/M2 macrophage activation

Upon injection of HP [1-^13^C]pyruvate into activated and control macrophages, HP [1-^13^C]lactate production could be detected at 183.3ppm, with the HP [1-^13^C]pyruvate resonance visible at 171.1ppm (**Figure 3A**). **Figure 3B** shows the mean HP [1-^13^C]lactate signal for control, M1 activated, and M2 activated datasets over the first 40 TRs normalized to cell number. The buildup of HP [1-^13^C]lactate can be observed, demonstrating *in situ* metabolism. Upon quantification of the total sum spectra, our results show that, in comparison to control cells, HP Lactate-to-Pyruvate ratio was significantly increased by 467±49% with M1 activation (p = 0.0010, 4.00±0.27 x10^-6^ for M1 activated cells *vs*. 7.06±0.39 x10^-7^ for Control cells). In M2 macrophages, a 91±36 % increase was observed compared to controls, but it did not reach significance (p = 0.1866, 1.35±0.24×10^-6^ for M2 activated cells) (**Figure 3C**). When comparing activation states, HP Lactate-to-Pyruvate ratio was significantly higher by 197±57% in M1 activated cells as compared to M2 activated cells (p = 0.0008).

**Figure 3.**
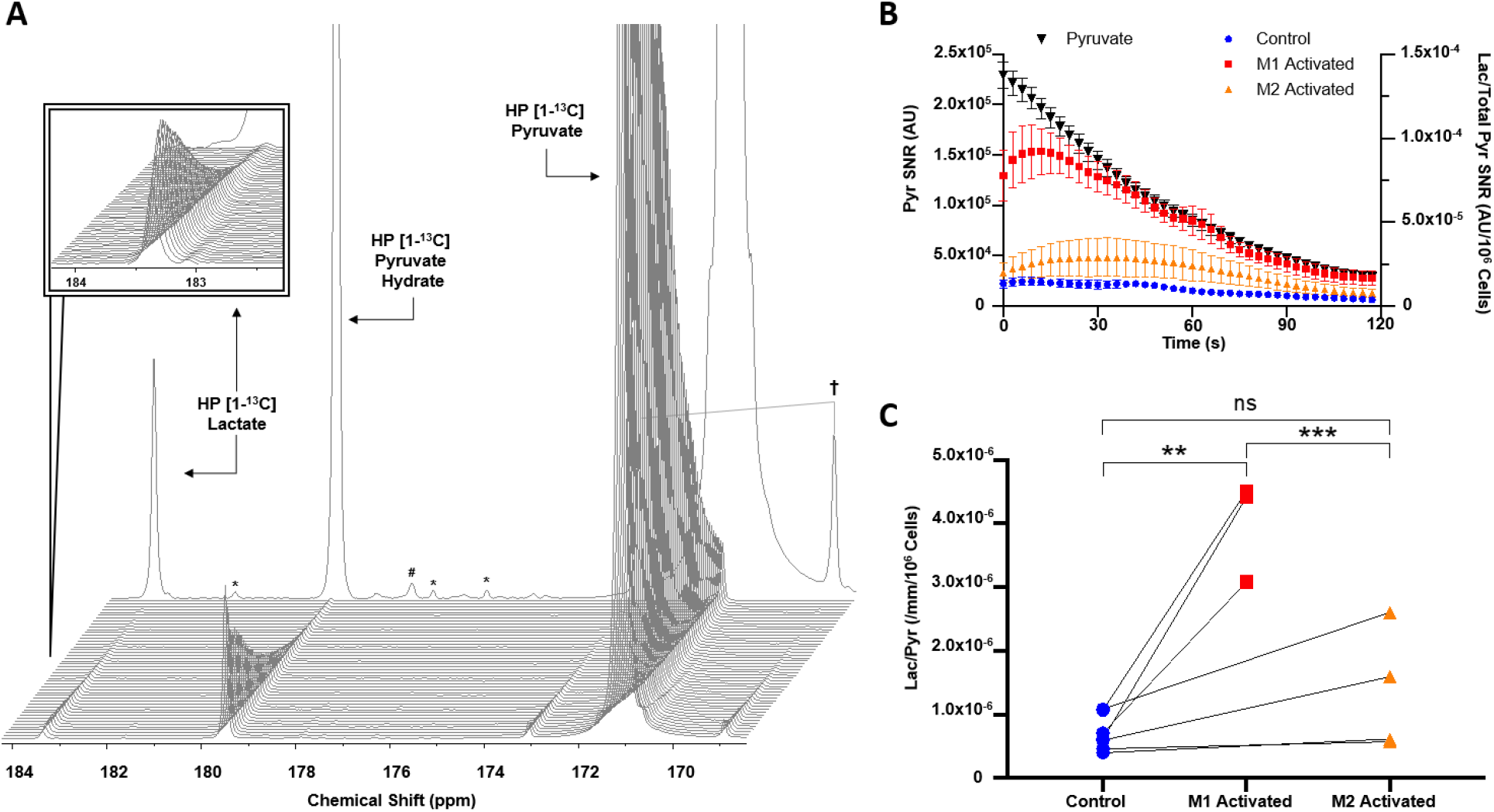
HP [1-^13^C]Lactate production is differentially increased by M1/M2 macrophage activation. (**A**) Dynamic spectra acquired at 1.47 Tesla post HP [1-^13^C]pyruvate injection into M1 activated J774a.1 macrophages, and corresponding sum spectra over 100 TRs in background. Insert shows HP [1-^13^C]lactate dynamics. († symmetric splitting of pyruvate due to 1.1% ^13^C natural abundance in C2 position; # alanine peak; * contaminants). (**B**) Signal to noise ratio (SNR) of HP [1-^13^C]pyruvate decay from all injections (black, left y-axis) and HP [1-^13^C]lactate production (right y-axis) in M1 activated (red), M2 activated (orange) and control cells (blue) over 120s. (**C**) HP lactate-to-pyruvate ratios measured in controls (blue) and their activation pairs (M1 activated red, M2 activated orange) (ns: p=0.1866, **p<0.01, ***p<0.001).

### HP [1-^13^C]DHA conversion to HP [1-^13^C]AA is increased with M1 activation

Following injection of HP [1-^13^C]DHA, HP [1-^13^C]AA production could be detected at 178.8ppm in control and activated macrophages, while the resonance of the substrate HP [1-^13^C]DHA was detected at 175ppm (**Figure 4A**). The time courses of HP [1-^13^C]DHA and HP [1-^13^C]AA are shown in **Figure 4B** for a control, an M1 and an M2 activated dataset. HP [1-^13^C]AA signal gradually builds up as the signal from HP [1-^13^C]DHA decays, denoting *in situ* conversion. Upon quantification, our results show that the HP AA-to-DHA ratio was significantly increased by 88±14 % in M1 activated cells compared to Control (p = 0.0034, 4.53±0.26 x10^-6^ in M1 activated *vs*. 2.41±0.11 x10^-6^ in Control), while no significant differences were observed in M2 activated macrophages *vs*. Control (p = 0.9581, 2.19±0.10 x10^-6^ in M2 activated) (**Figure 4C**). When comparing activation states, HP AA-to-DHA ratio was significantly higher by 107±15 % in M1 activated cells as compared to M2 activated cells (p = 0.0122).

**Figure 4.**
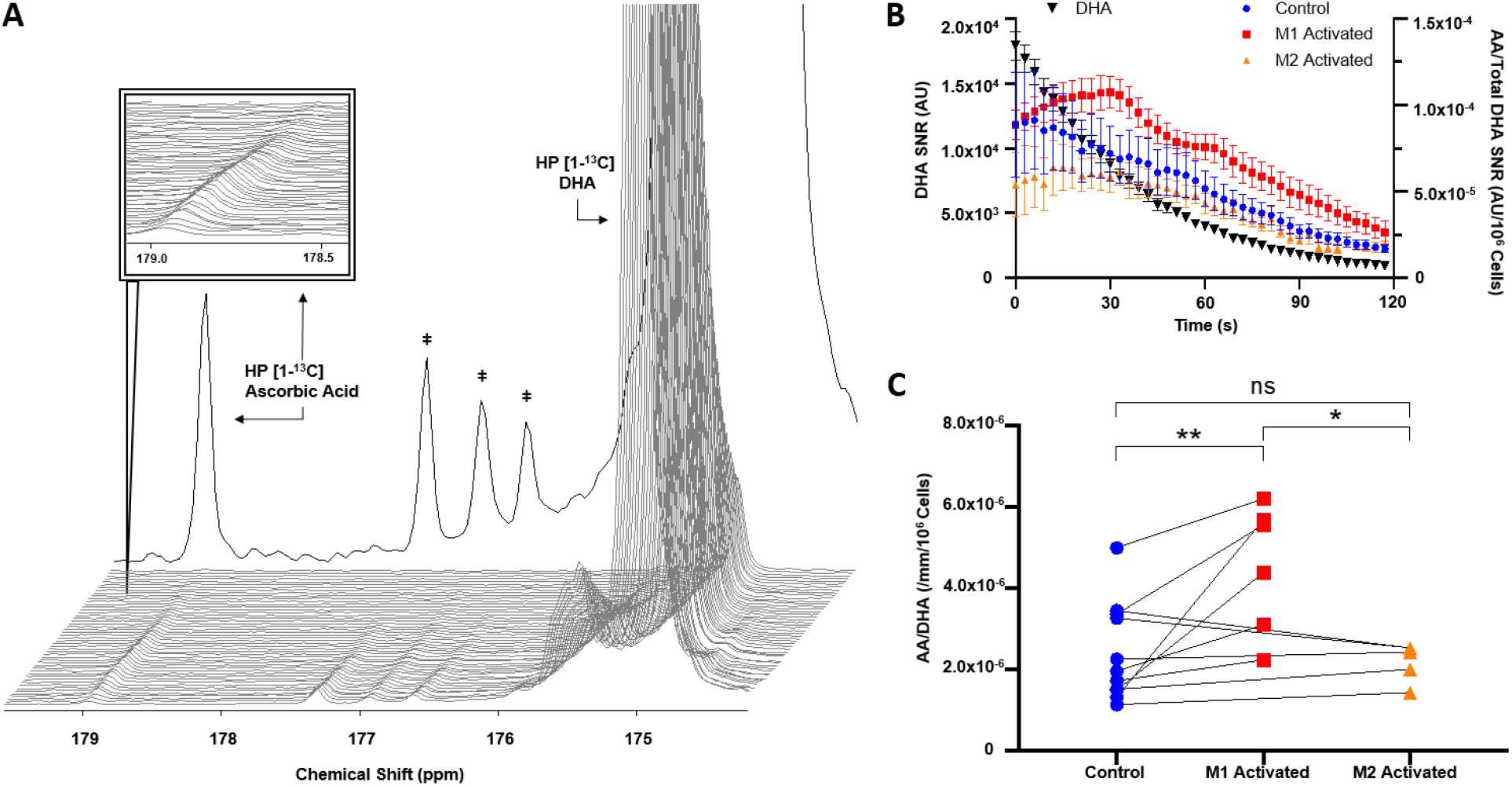
HP [1-^13^C]AA production is increased with M1 activation. (**A**) Dynamic spectra acquired at 1.47 Tesla post HP [1-^13^C]DHA injection into M1 activated J774a.1 macrophages, and sum spectra over 100 TRs in background (‡ contaminants). (**B**) Signal to noise ratio (SNR) of HP [1-^13^C]DHA decay from all injections (black, left y-axis) and HP [1-^13^C]AA production (right y-axis) in M1 activated (red), M2 activated (orange) and control cells (blue) over 120s. (**C**) HP AA-to-DHA ratios measured in controls (blue) and their activation pairs (M1 activated red, M2 activated orange) (ns: p=0.9581, *p<0.05, **p<0.01).

### LDH activity is increased in both M1 and M2 activated macrophages, while PDH activity is decreased in M1 and increased in M2 activation

Interestingly, LDH enzyme activity was significantly increased by 92±6% in M1 activated macrophages (p = 0.0020, 1.57×10^-4^ μM NADH/min/μg for M1 activated *vs*. 8.20×10^-5^ μM NADH/min/μg for Control), and by 27±6% in M2 activated macrophages (p = 0.0300, 1.04×10^-4^ μM NADH/min/μg) (**Figure 5A**). In contrast, PDH activity was significantly decreased by 34±5% in M1 activated macrophages (9.08×10^-4^ mOD/min/μg) compared to Control (1.37×10^-3^ mOD/min/μg, p = 0.0297). Meanwhile, M2 activated macrophages showed a significant 34±7% increase in PDH activity (1.83×10^-3^ mOD/min/μg) compared to Control (p = 0.0290) (**Figure 5B**). Spectrophotometric assay of GSH/GSSG ratios with subsequent normalization to 10^5^ cells showed no significant differences between groups (**Figure 5C**).

**Figure 5.**
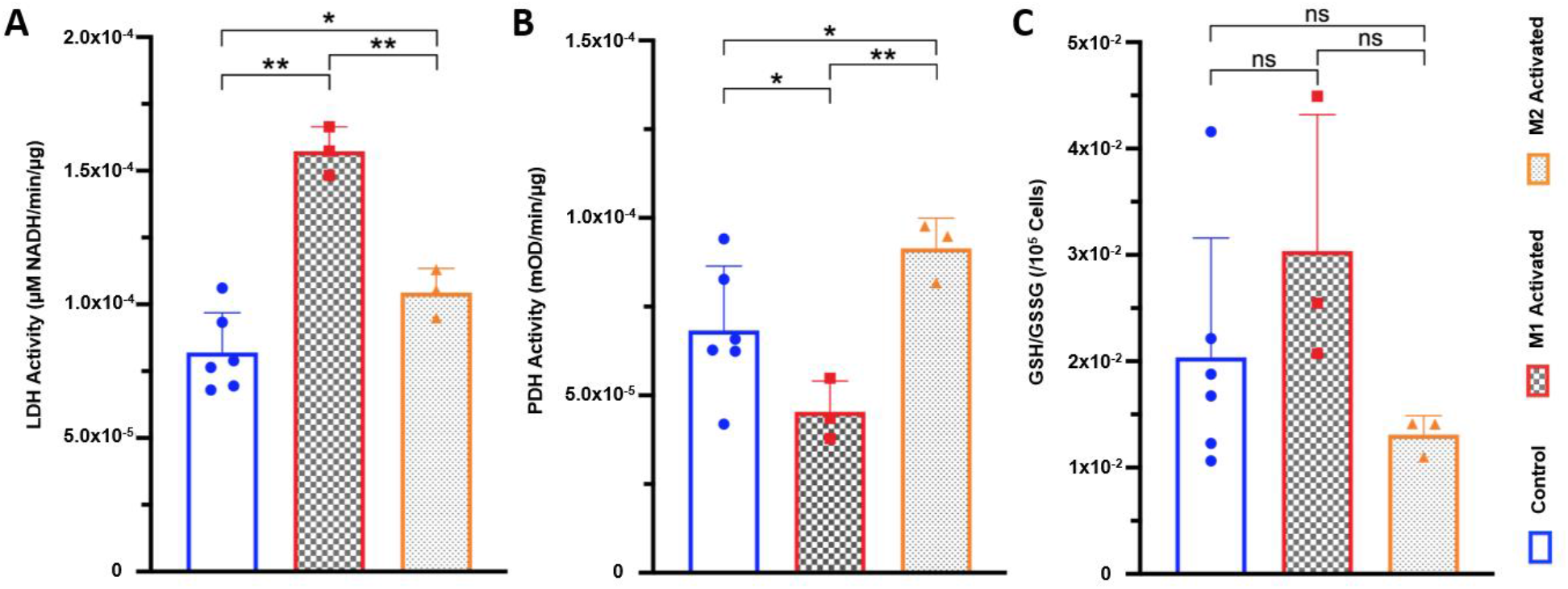
LDH and PDH activities are differentially modulated with M1/M2 activation. (**A**) Lactate dehydrogenase (LDH) enzyme activity for control (blue), M1 activated (red) and M2 activated (orange) macrophages (*p<0.05, **p<0.01, normalized to protein concentration). (**B**) Pyruvate dehydrogenase (PDH) enzyme activity measured by spectrophotometry for control (blue), M1 activated (red) and M2 activated (orange) macrophages (*p<0.05, **p<0.01, normalized to protein concentration). (**C**) Reduced glutathione (GSH) over oxidized glutathione (glutathione disulfide, GSSH) GSH/GSSG ratio per 10^5^ cells for control (blue), M1 activated (red) and M2 activated (orange) macrophages (ns control vs M1: p = 0.1265; ns control vs M2: p = 0.8326; ns M1 vs M2: p = 0.1083).

### High resolution ^1^H NMR detects significant differences in extracted metabolites between Control, M1, and M2 macrophages

^1^H NMR spectra of extracted metabolites exhibited well-resolved peaks at 800Mhz (**Figure 6A**). Multiple metabolite concentrations were observed to be significantly altered between groups (**Figure 6B**), with many in agreeance with previous reports^23^. In M1 activated macrophages, significant increases in itaconate (p<0.0001 vs. Control, p<0.0001 vs. M2), taurine (p<0.05 vs. Control, p<0.05 vs. M2) and succinate (p<0.001 vs. Control, p<0.0001 vs. M2) were measured, whereas a significant decrease in aspartate (p<0.05 vs. Control, p<0.05 vs. M2) was detected. In M2 macrophages, significant decreases in NAD (p<0.01 vs. Control, *Non Significant (NS)* vs. M1), lactate (p<0.05 vs. Control, p<0.05 vs. M1) and choline (p<0.05 vs. Control, *NS* vs. M1) were observed. Additional significant metabolite changes between M1 and M2 activated macrophages include glutamate (p<0.05), creatine (p<0.05), and arginine (p<0.05). Metabolite concentrations for each activation group are detailed in **Table 1**.

**Figure 6.**
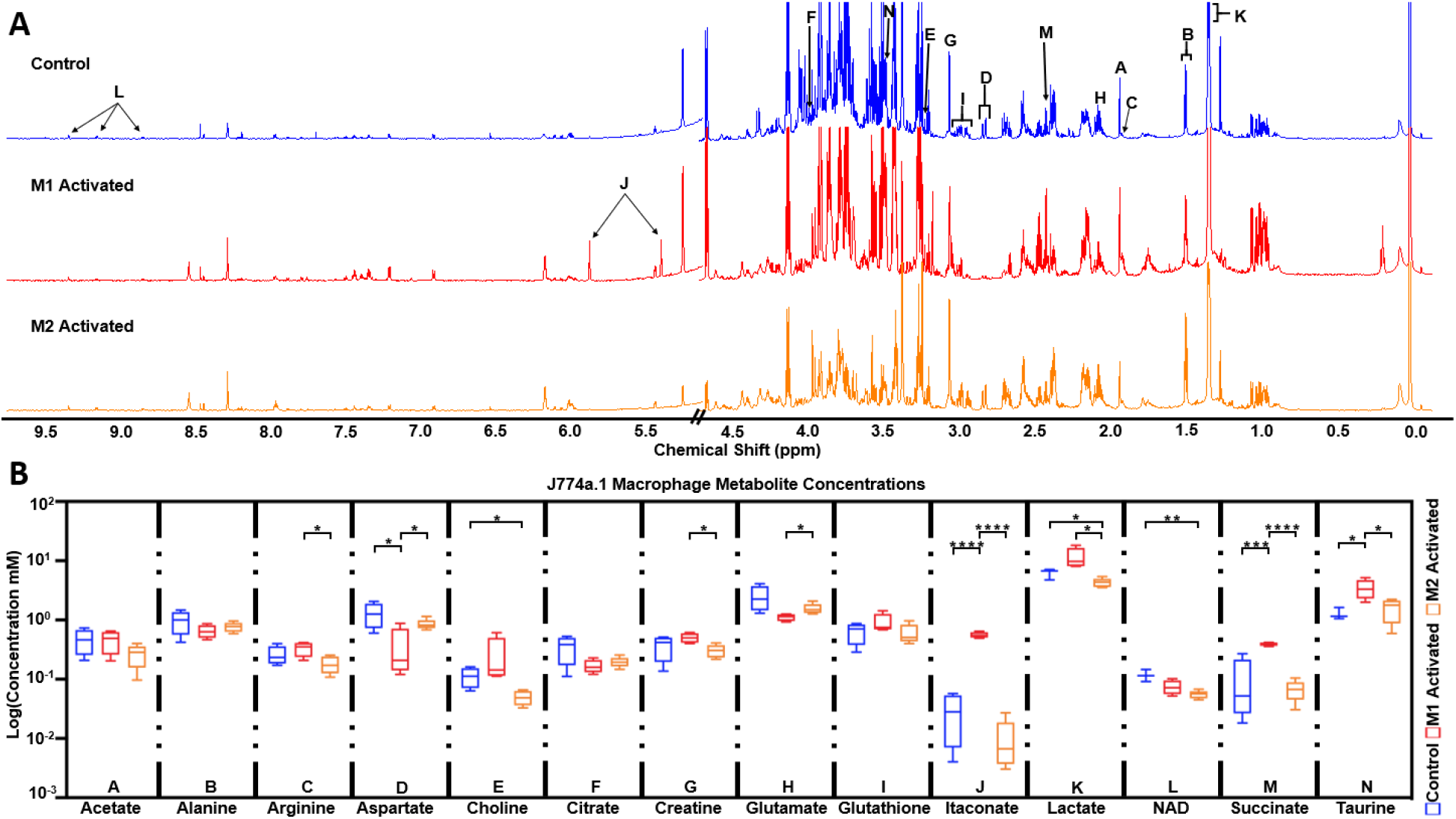
High resolution ^1^H NMR detects significant differences in extracted metabolites between Control, M1, and M2 macrophages. (**A**) Representative high resolution ^1^H NMR 1D NOESY spectra of control (blue), M1 activated (red), and M2 activated (orange) J774a.1 macrophage extracts (water region omitted; normalized to cell number with axes scaled identically). (**B**) Metabolite concentrations between control (blue; n = 4), M1 activated (red; n = 5), and M2 activated (orange; n = 5) determined from ^1^H NMR data. Data reported in log for purpose of presentation, with statistics performed on metabolite concentration (mM) normalized to cell number and TSP. Significance from two-way ANOVA shown between relevant groups with *p<0.05, **p<0.01, ***p<0.001, ****p<0.0001.

**Table 1.** Metabolite concentrations for control (blue), M1 activated (red), and M2 activated (orange) J774a.1 macrophages, calculated from high resolution 800MHz ^1^H NMR spectra of J774a.1 macrophage extracts. Concentration is reported as mean ± standard deviation.

| **J774a.1 Non-activated vs Activated Metabolite Concentrations (mM)** |  |  |  |
| --- | --- | --- | --- |
| **Metabolite** | **Control** | **M1 Activated** | **M2 Activated** |
| Acetate | 0.47 ± 0.22 | 0.46 ± 0.19 | 0.26 ± 0.11 |
| Alanine | 0.98 ± 0.43 | 0.65 ± 0.17 | 0.77 ± 0.14 |
| Arginine | 0.26 ± 0.10 | 0.33 ± 0.09 | 0.18 ± 0.06 |
| Aspartate | 1.29 ± 0.61 | 0.35 ± 0.35 | 0.85 ± 0.18 |
| Choline | 0.11 ± 0.04 | 0.25 ± 0.24 | 0.05 ± 0.01 |
| Citrate | 0.35 ± 0.18 | 0.17 ± 0.05 | 0.20 ± 0.04 |
| Creatine | 0.37 ± 0.17 | 0.50 ± 0.10 | 0.31 ± 0.08 |
| Glutamate | 2.48 ± 1.21 | 1.12 ± 0.15 | 1.56 ± 0.32 |
| Glutathione | 0.64 ± 0.25 | 0.90 ± 0.36 | 0.61 ± 0.23 |
| Itaconic acid | 0.03 ± 0.02 | 0.56 ± 0.06 | 0.01 ± 0.01 |
| Lactate | 6.21 ± 1.28 | 11.44 ± 4.74 | 4.29 ± 0.75 |
| NAD | 0.12 ± 0.03 | 0.07 ± 0.02 | 0.06 ± 0.01 |
| Succinate | 0.10 ± 0.12 | 0.40 ± 0.03 | 0.07 ± 0.03 |
| Taurine | 1.28 ± 0.31 | 3.45 ± 1.30 | 1.60 ± 0.70 |

## Discussion

In this study, we used the established J774a.1 mouse macrophage cell line and applied M1^23^ and M2^26^ activation protocols described in previous reports. First, we confirmed that differential activation was successfully achieved, as shown by a significant increase of arginase activity in M2 activated macrophages^30^ and an increased level of ROS in M1 activated macrophages^7^. Of note, we did not see a significant decrease in ROS levels in M2 activated macrophages compared to control, as others have reported^12^, likely due to the small number of cells collected in the subsamples and the subsequent lower sensitivity of the assay. Protein concentrations per cell were unchanged with M1 or M2 activation, justifying the normalization of all HP and spectrophotometric values to number of cells throughout the study. The ^1^H MRS results further confirmed differential activation of macrophages. Highly significant increases in itaconate and succinate concentrations are seen in M1 activated macrophages compared to control, in line with previous reports ^31^, as itaconate is a well-known product of M1 polarization^32^ and a potent inhibitor of succinate dehydrogenase which leads to succinate accumulation^33^. The increased lactate observed in M1 compared to control and M2 is also indicative of highly upregulated lactate production and Warburg-like effect that are specific to M1 activation^12^. In M2 activated macrophages, the observed decrease in arginine levels compared to M1 could be explained by increased arginine consumption through upregulated arginase activity, though to date the only metabolomics study of macrophage activation reports increased intracellular arginine concentration for both M1 and M2^34^. It should be noted however that this study used primary human macrophages, which have been shown to respond differently than established cell lines, as well as much longer activation protocol of 72 hours, which might explain the observed discrepancy^35^. A larger metabolomics study specific to activation patterns of murine macrophage cell lines would need to be pursued for more direct comparisons.

Here, we used an innovative approach allowing for noninvasive assessment of metabolism, namely HP ^13^C MRS, and evaluated its potential to differentiate between activation states in live macrophages. Our results show that significant differences in HP product-to-substrate ratios of both HP [1-^13^C]pyruvate and HP [1-^13^C]DHA can be observed in differentially activated murine macrophages. The sampling of cell slurries prior to each HP ^13^C MRS acquisition allowed for paired biochemical assays to be performed, thus enabling for strong biochemical validations.

The increased HP ^13^C lactate-to-pyruvate ratio observed in M1 activated macrophages is in agreeance with previous reports^23^, and congruent with the expected highly increased glycolytic activity in pro-inflammatory macrophages^36^. This increased ratio is also associated with a significant increase in LDH activity in the M1 group detected in paired subsamples. In M2 activated macrophages, the HP ^13^C lactate-to-pyruvate ratio was not significantly different from control, although a trend towards significance was observed. This trend is in line with a smaller significant increase in LDH activity, which was detected by spectrophotometric assays that are likely more sensitive than HP ^13^C MRS. These results are also in agreement with previous reports showing that M2 polarized macrophages exhibit a more modest increase in glycolytic activity compared to non-activated macrophages^37^. To further understand our HP results, it is important to look into another common pathway for HP [1-^13^C]pyruvate metabolism, through PDH. In M1 activated macrophages, our spectrophotometric results showing decreased PDH activity compared to controls are in line with previous reports showing significantly reduced PDH activity in M1 primary murine bone marrow derived macrophages^31^. Reduced PDH activity in M1 macrophages would lead to less HP [1-^13^C]pyruvate entering the Krebs cycle, shuttling this HP probe towards LDH and increased HP [1-^13^C]lactate production, in line with our HP results. In M2 activated macrophages on the other hand, we observed a significant increase in PDH activity. This result is interesting, and reasonable given that expression of pyruvate dehydrogenase kinase (PDK), an inhibitor of PDH, is reduced in M2 polarization^38^. In that case, and contrary to M1, increased PDH activity may lead to more HP [1-^13^C]pyruvate being shuttled into the mitochondria, thus decreasing flux towards HP [1-^13^C]lactate production via LDH. This result provides an additional explanation for the fact that HP lactate-pyruvate ratio was not significantly increased in M2 activated macrophages, despite the increased LDH activity. Finally, in addition to enzymatic activities, the levels of steady-state lactate levels from ^1^H NMR might also contribute to the HP readouts through the well-documented pool-size effect^39-41^. The observed significant decrease in ^1^H lactate levels in M2 macrophages may also contribute to the reduced HP lactate levels in that group. In M1 macrophages, lactate levels follow an increasing trend (p = 0.128), which is also in line with the HP results. Multiple additional factors contributing to the HP readouts could be considered, including levels of membrane transporters (e.g. MCT1) or NAD cofactor availability. However, such measures could not be performed in this study due to the limited number of paired samples available.

Our results show that conversion of HP [1-^13^C]DHA to HP [1-^13^C]AA was increased in M1 activated macrophages, mirrored by increased ROS levels, but unexpectedly not by any changes in GSH levels or GSH/GSSG ratio. It is well known that the GSH redox system is a mitigator of ROS^42^, while also being coupled to the conversion of DHA into AA^43^: GSH is oxidized to GSSG by DHA, which in turn is converted to AA. Increasing ROS levels should theoretically deplete the available pool of GSH (and increase levels of GSSG), leaving less GSH available for the production of AA, and thus leading to a decreased HP AA-to-DHA ratio, which has been previously reported^25,44^. However, both GSH/GSSG ratios, as detected by spectrophotometric assay, and total GSH levels, as detected by ^1^H NMR, were not significantly different between M1 macrophages and control or M2. A previous study of the same cell line also showed that the total pool of GSH as detected by ^1^H NMR was not increased post activation by LPS^23^. Other reports using biochemical methods also show no differences in total GSH levels with M1 activation, but do note elevated GSH/GSSG ratio^45^. Elevated GSH/GSSG ratio can explain an increase in AA production, and may be a compensatory effect unique to macrophages as they endogenously manufacture ROS as part of the immune response. Our results show that a trend to an increase in M1 activated compared to control (p = 0.1265) and M2 (p = 0.1083) was seen, although it did not reach significance possibly due to the limited numbers of available paired samples (n = 3) that were spread thinly across the numerous assays. Further work could be done with a larger sample size to confirm the increase in GSH/GSSG ratio in M1 activated macrophages.

The power of dynamic metabolic probing using HP ^13^C MR is compelling, as it allows measuring previously inaccessible metabolic reactions non-invasively. HP ^13^C metabolic imaging is a rapidly growing field, and is now being used in multiple clinical trials across the globe, targeting several organs including brain, heart, prostate and kidney^46-57^. It is very likely that, given the constant improvements reported both on the acquisition and processing sides, this methodology will soon approach feasibility for widespread clinical adoption. Currently, the most popular probe for hyperpolarization is HP [1-^13^C]pyruvate, with some of the best polarization characteristics^41^, but other probes are continuously being investigated, opening up possibilities not explored before. While the clinical data reported so far is highly compelling, mechanistic studies are still critically needed to understand the relative contribution of each cell type to the detected HP signal, especially for cells as ubiquitous as macrophages, which are found in most diseases and most organs. Here, we performed the first study of live macrophages at a clinically-relevant field strength, and compared both M1 and M2 activation patterns. Before, the only other study that used HP ^3^C MRS on activated macrophages was conducted at the high magnetic field strength of 11.7 Tesla, and employed M1 activation only^23^. We showed that, at clinical field strength, the signal to noise ratio of substrate and products was sufficient to enable measurements of metabolic fluxes in live cells. Further, we demonstrated that M1 and M2 macrophages have a different HP metabolic signature, with both HP [1-^13^C]pyruvate and HP [1-^13^C]DHA. This highly significant difference should help our understanding of the metabolic readouts observed *in vivo* in preclinical models and patients.

## Acknowledgements

This work was supported by research grants: NIH R01NS102156, NMSS research grant RG-1701-26630, Hilton Foundation – Marilyn Hilton Award for Innovation in MS Research #17319. Dana Foundation: The David Mahoney Neuroimaging program and the NIH Hyperpolarized MRI Technology Resource Center #P41EB013598.

## Author contributions

KQ and LLP collected data. KQ performed the data analysis. MMC conceptualized and designed the overall study. All authors contributed equally to the text and reviewed the manuscript.

## Notes

### Competing Interest Statement

The authors have declared no competing interest.

## References

1. Galli, S. J., Borregaard, N. & Wynn, T. A. Phenotypic and functional plasticity of cells of innate immunity: macrophages, mast cells and neutrophils. Nature immunology 12, 1035–1044, doi:10.1038/ni.2109 (2011).

2. Okabe, Y. & Medzhitov, R. Tissue biology perspective on macrophages. Nat Immunol 17, 9–17, doi:10.1038/ni.3320 (2016).

3. Gasteiger, G. et al. Cellular Innate Immunity: An Old Game with New Players. Journal of Innate Immunity 9, 111–125, doi:10.1159/000453397 (2017).

4. Shiomi, A. & Usui, T. Pivotal roles of GM-CSF in autoimmunity and inflammation. Mediators of inflammation 2015, 568543–568543, doi:10.1155/2015/568543 (2015).

5. Kraakman, M. J., Murphy, A. J., Jandeleit-Dahm, K. & Kammoun, H. L. Macrophage polarization in obesity and type 2 diabetes: weighing down our understanding of macrophage function? Frontiers in immunology 5, 470–470, doi:10.3389/fimmu.2014.00470 (2014).

6. Dandekar, R., Kingaonkar, A. & Dhabekar, G. Role of macrophages in malignancy. Annals of Maxillofacial Surgery 1, 150–154, doi:10.4103/2231-0746.92782 (2011).

7. Mills, C. D., Kincaid, K., Alt, J. M., Heilman, M. J. & Hill, A. M. M-1/M-2 macrophages and the Th1/Th2 paradigm. J Immunol 164, 6166–6173, doi:10.4049/jimmunol.164.12.6166 (2000).

8. Stunault, M. I., Bories, G., Guinamard, R. R. & Ivanov, S. Metabolism Plays a Key Role during Macrophage Activation. Mediators of Inflammation 2018, 2426138, doi:10.1155/2018/2426138 (2018).

9. Tan, H.-Y. et al. The Reactive Oxygen Species in Macrophage Polarization: Reflecting Its Dual Role in Progression and Treatment of Human Diseases. Oxidative Medicine and Cellular Longevity 2016, 2795090, doi:10.1155/2016/2795090 (2016).

10. Rendra, E. et al. Reactive oxygen species (ROS) in macrophage activation and function in diabetes. Immunobiology 224, 242–253, doi:10.1016/j.imbio.2018.11.010 (2019).

11. Yang, Z. & Ming, X.-F. Functions of arginase isoforms in macrophage inflammatory responses: impact on cardiovascular diseases and metabolic disorders. Frontiers in immunology 5, 533–533, doi:10.3389/fimmu.2014.00533 (2014).

12. Martinez, F. O. & Gordon, S. The M1 and M2 paradigm of macrophage activation: time for reassessment. F1000prime reports 6, 13–13, doi:10.12703/P6-13 (2014).

13. Ardenkjaer-Larsen, J. H. et al. Increase in signal-to-noise ratio of > 10,000 times in liquid-state NMR. Proceedings of the National Academy of Sciences of the United States of America 100, 10158–10163, doi:10.1073/pnas.1733835100 (2003).

14. Chaumeil, M. M. et al. Non-invasive in vivo assessment of IDH1 mutational status in glioma. Nature Communications 4, 2429, doi:10.1038/ncomms3429 (2013).

15. Le Page, L. M., Guglielmetti, C., Taglang, C. & Chaumeil, M. M. Imaging Brain Metabolism Using Hyperpolarized ^13^C Magnetic Resonance Spectroscopy. Trends in Neurosciences 43, 343–354, doi:10.1016/j.tins.2020.03.006 (2020).

16. Yoshihara, H. A., Bastiaansen, J. A., Berthonneche, C., Comment, A. & Schwitter, J. An intact small animal model of myocardial ischemia-reperfusion: Characterization of metabolic changes by hyperpolarized 13C MR spectroscopy. Am J Physiol Heart Circ Physiol 309, H2058–2066, doi:10.1152/ajpheart.00376.2015 (2015).

17. Kurhanewicz, J. et al. Hyperpolarized (13)C MRI: Path to Clinical Translation in Oncology. Neoplasia (New York, N.Y.) 21, 1–16, doi:10.1016/j.neo.2018.09.006 (2019).

18. ClinicalTrials.gov. Imaging of Traumatic Brain Injury Metabolism Using Hyperpolarized Carbon-13 Pyruvate. National Library of Medicine (US) (2018).

19. Nelson, S. J. et al. Metabolic imaging of patients with prostate cancer using hyperpolarized [1-^13^C]pyruvate. Science translational medicine 5, 198ra108–198ra108, doi:10.1126/scitranslmed.3006070 (2013).

20. Le Page, L. M., Guglielmetti, C., Taglang, C. & Chaumeil, M. M. Imaging Brain Metabolism Using Hyperpolarized (13)C Magnetic Resonance Spectroscopy. Trends Neurosci 43, 343–354, doi:10.1016/j.tins.2020.03.006 (2020).

21. Grist, J. T. et al. Hyperpolarized (13)C MRI: A novel approach for probing cerebral metabolism in health and neurological disease. J Cereb Blood Flow Metab 40, 1137–1147, doi:10.1177/0271678X20909045 (2020).

22. Najac, C. et al. Detection of inflammatory cell function using 13C magnetic resonance spectroscopy of hyperpolarized [6-13C]-arginine. Scientific Reports 6, 31397, doi:10.1038/srep31397 (2016).

23. Sriram, R. et al. Molecular detection of inflammation in cell models using hyperpolarized (13)C-pyruvate. Theranostics 8, 3400–3407, doi:10.7150/thno.24322 (2018).

24. Kohler, S. J. et al. In vivo 13 carbon metabolic imaging at 3T with hyperpolarized 13C-1-pyruvate. Magn Reson Med 58, 65–69, doi:10.1002/mrm.21253 (2007).

25. Keshari, K. R. et al. Hyperpolarized 13C dehydroascorbate as an endogenous redox sensor for in vivo metabolic imaging. Proceedings of the National Academy of Sciences of the United States of America 108, 18606–18611, doi:10.1073/pnas.1106920108 (2011).

26. Csóka, B. et al. Adenosine promotes alternative macrophage activation via A2A and A2B receptors. FASEB journal: official publication of the Federation of American Societies for Experimental Biology 26, 376–386, doi:10.1096/fj.11-190934 (2012).

27. Sapcariu, S. C. et al. Simultaneous extraction of proteins and metabolites from cells in culture. MethodsX 1, 74–80, doi:10.1016/j.mex.2014.07.002 (2014).

28. Wishart, D. S. et al. HMDB 4.0: the human metabolome database for 2018. Nucleic Acids Res 46, D608–d617, doi:10.1093/nar/gkx1089 (2018).

29. Koo, S.-J., Chowdhury, I. H., Szczesny, B., Wan, X. & Garg, N. J. Macrophages Promote Oxidative Metabolism To Drive Nitric Oxide Generation in Response to Trypanosoma cruzi. Infection and immunity 84, 3527–3541, doi:10.1128/IAI.00809-16 (2016).

30. Rath, M., Müller, I., Kropf, P., Closs, E. I. & Munder, M. Metabolism via Arginase or Nitric Oxide Synthase: Two Competing Arginine Pathways in Macrophages. Frontiers in immunology 5, 532–532, doi:10.3389/fimmu.2014.00532 (2014).

31. Palmieri, E. M. et al. Nitric oxide orchestrates metabolic rewiring in M1 macrophages by targeting aconitase 2 and pyruvate dehydrogenase. Nature Communications 11, 698, doi:10.1038/s41467-020-14433-7 (2020).

32. Michelucci, A. et al. Immune-responsive gene 1 protein links metabolism to immunity by catalyzing itaconic acid production. Proceedings of the National Academy of Sciences 110, 7820, doi:10.1073/pnas.1218599110 (2013).

33. Cordes, T. et al. Immunoresponsive Gene 1 and Itaconate Inhibit Succinate Dehydrogenase to Modulate Intracellular Succinate Levels. J Biol Chem 291, 14274–14284, doi:10.1074/jbc.M115.685792 (2016).

34. Fuchs, A. L. et al. Quantitative (1)H NMR Metabolomics Reveal Distinct Metabolic Adaptations in Human Macrophages Following Differential Activation. Metabolites 9, 248, doi:10.3390/metabo9110248 (2019).

35. Andreu, N. et al. Primary macrophages and J774 cells respond differently to infection with Mycobacterium tuberculosis. Scientific reports 7, 42225–42225, doi:10.1038/srep42225 (2017).

36. Hu, S. et al. 13C-pyruvate imaging reveals alterations in glycolysis that precede c-Myc-induced tumor formation and regression. Cell metabolism 14, 131–142, doi:10.1016/j.cmet.2011.04.012 (2011).

37. Huang, S. C.-C. et al. Metabolic Reprogramming Mediated by the mTORC2-IRF4 Signaling Axis Is Essential for Macrophage Alternative Activation. Immunity 45, 817–830, doi:10.1016/j.immuni.2016.09.016 (2016).

38. Min, B.-K. et al. Pyruvate Dehydrogenase Kinase Is a Metabolic Checkpoint for Polarization of Macrophages to the M1 Phenotype. Frontiers in immunology 10, 944–944, doi:10.3389/fimmu.2019.00944 (2019).

39. Hong, C. S. et al. MCT1 Modulates Cancer Cell Pyruvate Export and Growth of Tumors that Co-express MCT1 and MCT4. Cell reports 14, 1590–1601, doi:10.1016/j.celrep.2016.01.057 (2016).

40. Vincent, S. J., Zwahlen, C., Post, C. B., Burgner, J. W. & Bodenhausen, G. The conformation of NAD+ bound to lactate dehydrogenase determined by nuclear magnetic resonance with suppression of spin diffusion. Proceedings of the National Academy of Sciences of the United States of America 94, 4383–4388, doi:10.1073/pnas.94.9.4383 (1997).

41. Chaumeil, M. M., Najac, C. & Ronen, S. M. Studies of Metabolism Using (13)C MRS of Hyperpolarized Probes. Methods Enzymol 561, 1–71, doi:10.1016/bs.mie.2015.04.001 (2015).

42. Armstrong, J. S. et al. Role of glutathione depletion and reactive oxygen species generation in apoptotic signaling in a human B lymphoma cell line. Cell Death & Differentiation 9, 252–263, doi:10.1038/sj.cdd.4400959 (2002).

43. Linster, C. L. & Van Schaftingen, E. Vitamin C. Biosynthesis, recycling and degradation in mammals. Febs j 274, 1–22, doi:10.1111/j.1742-4658.2006.05607.x (2007).

44. Qin, H. et al. Imaging glutathione depletion in the rat brain using ascorbate-derived hyperpolarized MR and PET probes. Scientific reports 8, 7928–7928, doi:10.1038/s41598-018-26296-6 (2018).

45. Kadl, A. et al. Identification of a novel macrophage phenotype that develops in response to atherogenic phospholipids via Nrf2. Circ Res 107, 737–746, doi:10.1161/circresaha.109.215715 (2010).

46. Autry, A. W. et al. Comparison between 8- and 32-channel phased-array receive coils for in vivo hyperpolarized (13) C imaging of the human brain. Magn Reson Med 82, 833–841, doi:10.1002/mrm.27743 (2019).

47. Chung, B. T. et al. First hyperpolarized [2-(13)C]pyruvate MR studies of human brain metabolism. J Magn Reson 309, 106617, doi:10.1016/j.jmr.2019.106617 (2019).

48. Gordon, J. W. et al. Translation of Carbon-13 EPI for hyperpolarized MR molecular imaging of prostate and brain cancer patients. Magn Reson Med 81, 2702–2709, doi:10.1002/mrm.27549 (2019).

49. Nelson, S. J. et al. Metabolic imaging of patients with prostate cancer using hyperpolarized [1-(1)(3)C]pyruvate. Sci Transl Med 5, 198ra108, doi:10.1126/scitranslmed.3006070 (2013).

50. Lee, C. Y. et al. Lactate topography of the human brain using hyperpolarized (13)C-MRI. Neuroimage 204, 116202, doi:10.1016/j.neuroimage.2019.116202 (2019).

51. Miloushev, V. Z. et al. Metabolic Imaging of the Human Brain with Hyperpolarized (13)C Pyruvate Demonstrates (13)C Lactate Production in Brain Tumor Patients. Cancer Res 78, 3755–3760, doi:10.1158/0008-5472.CAN-18-0221 (2018).

52. Park, I. et al. Development of methods and feasibility of using hyperpolarized carbon-13 imaging data for evaluating brain metabolism in patient studies. Magn Reson Med 80, 864–873, doi:10.1002/mrm.27077 (2018).

53. Park, J. M. Imaging of Traumatic Brain Injury Metabolism Using Hyperpolarized Carbon-13 Pyruvate - ClinicalTrials.gov, <https://clinicaltrials.gov/ct2/show/NCT03502967> (2019).

54. Gallagher, F. A. et al. Imaging breast cancer using hyperpolarized carbon-13 MRI. Proc Natl Acad Sci U S A 117, 2092–2098, doi:10.1073/pnas.1913841117 (2020).

55. Granlund, K. L. et al. Hyperpolarized MRI of Human Prostate Cancer Reveals Increased Lactate with Tumor Grade Driven by Monocarboxylate Transporter 1. Cell Metab 31, 105–114 e103, doi:10.1016/j.cmet.2019.08.024 (2020).

56. Grist, J. T. et al. Creating a clinical platform for carbon-13 studies using the sodium-23 and proton resonances. Magn Reson Med 84, 1817–1827, doi:10.1002/mrm.28238 (2020).

57. Woitek, R. et al. Hyperpolarized (13)C MRI of Tumor Metabolism Demonstrates Early Metabolic Response to Neoadjuvant Chemotherapy in Breast Cancer. Radiol Imaging Cancer 2, e200017, doi:10.1148/rycan.2020200017 (2020).

